# Role of horizontal gene transfers and microbial ecology in the evolution of fluxes through the tricarboxylic acid cycle

**DOI:** 10.1101/2022.10.25.513756

**Authors:** Tymofii Sokolskyi, Shiladitya DasSarma

## Abstract

The origin of carbon fixation is a fundamental question in astrobiology. While the Calvin cycle is the most active on the modern Earth, the reductive TCA cycle (rTCA) pathway for carbon fixation has been proposed to have played an important role in early evolution. In this study, we examined the evolution of key enzymes in the rTCA, which are rare in extant organisms, occurring in a few groups of Bacteria and Archaea. We investigated one of the least common reactions of this pathway, cleavage of citrate into oxaloacetate and acetyl-CoA, which can be performed by either a two-enzyme system (CCS/CCL) or a single enzyme (ACL) that is assumed to be the result of fusion of the two active sites into a single polypeptide. For broader context, we also studied functionally diverged homologs of these enzymes, succinyl-CoA synthetase (SCS) and citrate synthase (CS). Our phylogenetic analysis of these enzymes in Bacteria and Archaea shows that SCS, a homolog of CCS from distant bacterial taxa capable of citrate cleavage, are monophyletic, suggesting linked horizontal gene transfers of SCS and citrate cleavage enzymes. We also found evidence of the horizontal transfer of SCS from a clade of anaerobic Archaea (Archaeoglobi, Methanomicrobia or Crenarchaeota) to an ancestor of Cyanobacteria/Melainabacteria clade – both of whom share a succinate semialdehyde shunt in their oxidative TCA cycles. We also identified new bacterial and archaeal taxa for which complete rTCA cycles are theoretically possible, including *Syntrophobacter, Desulfofundulus, Beggiatoa, Caldithrix, Ca*. Acidulodesulfobacterales and *Ca*. Micrarachaeota. Finally, we suggest a possibility for syntrophically-regulated fluxes through oxidative and reductive TCA reactions in microbial communities particularly Haloarchaea-Nanohaloarchaea symbiosis and its implications for the Purple Earth hypothesis. We discuss how the inclusion of an ecological perspective in the studies of evolution of ancient metabolic pathways may be beneficial to understand origins of life.

## Introduction

The tricarboxylic acid (TCA) cycle is a well-known metabolic pathway, widely distributed among aerobic organisms that produces reducing equivalents and is an integral part of aerobic respiration. In most organisms the TCA runs exclusively in the oxidative direction. However, it was long known that some of the enzymes of the oxidative TCA cycle (oTCA) can catalyze their respective reverse reactions. In 1966, Evans et al. observed the presence of a unique carbon fixation pathway in a photosynthetic bacterium *Chlorobium thiosulfatophilum* that results in incorporation of four molecules of carbon dioxide during a full cycle. This cycle, reductive TCA (rTCA), shares most of its reactions with oTCA, except three reactions that are catalyzed by different enzymes. These rTCA-specific reactions are: (i) citrate cleavage into oxaloacetate and acetyl-CoA, catalyzed by ATP-citrate lyase (ACL); (ii) fumarate-succinate conversion by fumarate reductase (FR); and (iii) succinyl-CoA to 2-oxoglutarate by ferredoxin-dependent 2-oxoglutarate synthase (FOS) (Buchanan et al., 2017). Of these reactions, citrate cleavage is particularly interesting since it is the branching reaction of the autocatalytic cycle, as defined by Peng et al., 2020, that produces two oxaloacetate molecules per round (Fig. 1, A).

**Fig. 1.**
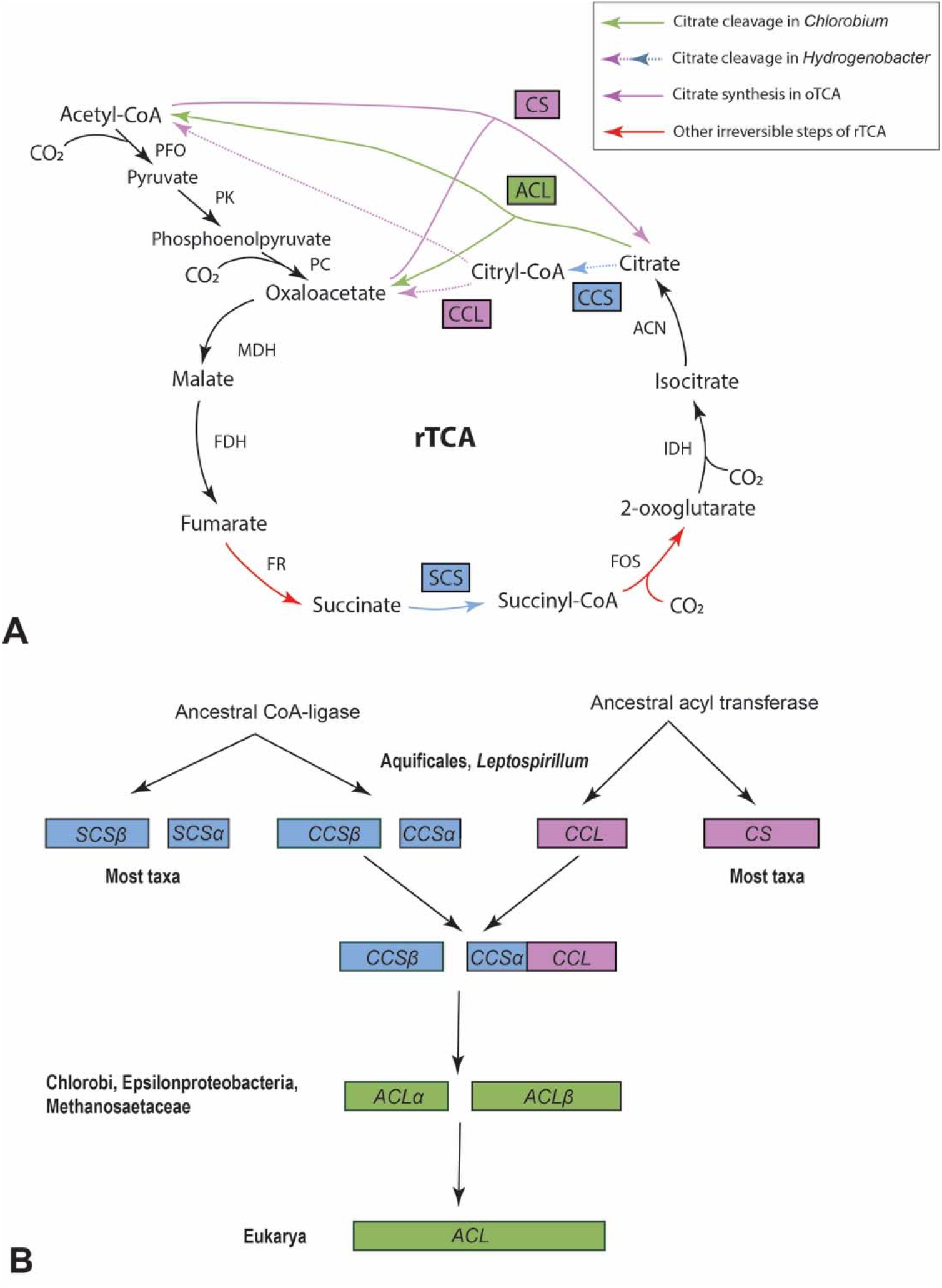
Overview of the rTCA and citrate cleavage in prokaryotes. A. Scheme of the reductive TCA cycle illustrating two known mechanisms of citrate cleavage into oxaloacetate and acetyl-CoA: two enzyme CCS/CCL system as in *Aquifex* and one-step ACL reaction in *Chlorobium* (Berg, 2011). Examples of enzymes capable of catalyzing these reactions are noted above the arrows. B. Scheme of evolution of CCS, CCL, ACL, SCS and CS illustrating a sequence of fusions as proposed by Aoshima, 2007 and expanded on by our analyses. Abbreviations: CCS – citryl-CoA synthase; CCL – citryl-CoA lyase; ACL – ATP citrate lyase; PFO – pyruvate ferredoxin oxidoreductase; PK – pyruvate kinase; PC – phosphoenolpyruvate carboxylase; MDH – malate dehydrogenase; FDH – fumarate dehydratase; FR – fumarate reductase; FOS – ferredoxin 2-oxoglutarate synthase; IDH – isocitrate dehydrogenase; ACN - aconitase.

After its discovery (Evans et al. 1966), the rTCA remained poorly studied and known only from a handful of bacterial clades. It was described from anoxygenic green sulfur bacteria, such as *Chlorobium* (Tang & Blankenship, 2010), epsilon, delta and alpha Proteobacteria (Hugler et al., 2005; Williams et al., 2006; Schauder et al., 1987), *Nitrospira* (Bar-Even et al., 2012), and in bacteria from the phylum Aquificae (Hugler et al., 2007). The latter group is especially interesting, since one of the key steps of rTCA – citrate cleavage, is performed differently than in *Chlorobium* (Aoshima, 2004). As summarized in Fig. 1A, *Chlorobium* uses a larger enzyme, ATP citrate lyase (ACL; EC 2.3.3.8), which is composed of two subunits, to catalyze the citrate cleavage into oxaloacetate and acetyl-CoA (Aoshima, 2007). The citryl-CoA intermediate never leaves the ACL enzyme during this reaction. Aquificae, in contrast, have two separate enzymes: citryl-CoA synthetase (CCS; EC 6.2.1.18), which produces citryl-CoA from citrate, and citryl-CoA lyase (CCL; EC 4.1.3.34), which cleaves citryl-CoA (Aoshima et al., 2004). CCL and CCS were shown to be homologous to different subunits of ACL (Aoshima, 2007), implying that the bacterial ACL enzyme evolved by gene fusion with CCS large subunit being equivalent to ACL small subunit and CCS small subunit and CCL fusing into ACL large subunit (Fig. 1, B).

ACL, CCS and CCL proteins are known to share some homology to two other TCA enzymes – succinyl-CoA synthetase (SCS) and citrate synthase (CS). According to Aoshima (2007) and Beccerra et al. (2014), the SCS small subunit shares regions of homology to the CCS small subunit and part of the ACL large subunit. Similarly, the SCS large subunit is homologous to the CCS large and ACL small subunits and CS shares homology to CCL and part of the ACL large subunit. These homologies indicate a possibility of common descent for at least three rTCA enzymes, as illustrated in Fig. 1B. Taxonomic surveys suggest that citrate cleavage is by far the least common one among prokaryotes with the incomplete cycle (Aoshima, 2007). Most eukaryotes, on the other hand, possess an ATP-citrate lyase enzyme used to synthetize acetyl-CoA from citrate, even while there is no evidence of rTCA operating in this group (Chypre et al., 2012).

Carbon fixation pathways often operate in coordination with phototrophic pathways, as evident in Cyanobacteria with the Calvin-Benson cycle or Chlorobi with rTCA, due to the significant reduced electron carrier requirements of these cycles (Bar-Even et al., 2012). It was proposed earlier that retinal-based phototrophy, common in various groups of microbes such as Haloarchaea, could have been significantly more common prior to emergence of chlorophyll-based photosynthesis. Purple Earth hypothesis proposes that the spectral characteristics of chlorophyll-based photosystems might have resulted due to competition for wavelengths with Haloarchaea or other organisms using bacteriorhodopsin-like proteins (DasSarma & Schwieterman, 2021). Bacteriorhodopsins are light-activated proton pumps that were shown to be ancient and spectrally tuned to absorbing green light (Sephus et al., 2021). However, unlike most modern phototrophs that possess mechanisms for carbon fixation, extant Haloarchaea lack any carbon fixation pathways. Indeed, complete rTCA cycle specifically is currently unknown in any Archaea and most autotrophic members of this group use Wood-Ljungdahl pathway for fixing atmospheric carbon dioxide (Berg et al., 2010). Crenarchaeote *Thermoproteus neutrophilus* was described as capable of citrate cleavage and the entire rTCA pathway (Beh et al., 1993); however, this was later disproven (Becerra et al., 2014). In most Archaea, many of the key rTCA enzymes are missing and therefore, Archaea are frequently overlooked in the studies of rTCA evolution. However, Haloarchaea possess all but one – enzyme for citrate cleavage. This is one of the most complete rTCA pathways in any archaeon.

Yet, despite being largely absent from Archaea, multiple studies suggest ancient origins of rTCA, implying it might even be a primordial carbon fixation pathway in early organisms (Nunoura et al., 2018; Muchowska et al., 2017). This has gained support by evidence that many rTCA reactions can occur non-enzymatically (Zubarev et al., 2015), leading many to suspect that the Last Universal Common Ancestor (LUCA) was capable of autotrophic carbon fixation via the rTCA (Wimmer et al., 2021; Sutherland, 2017). Since the rTCA shows stoichiometric autocatalysis, producing 2 oxaloacetate molecules per cycle, and given that autocatalytic cycles might have allowed primordial heritability and responses to selection (Peng et al. 2022), the rTCA has been implicated as playing a key role in the origin of life (Wachtershauser, 1988; Smith & Morowitz 2016).

Despite the centrality of rTCA in theories of life’s origin, the scattered distribution of rTCA among modern organisms presents a problem. Here we used a large-scale phylogenetic analysis of ACL, CCS, CCL and their homologs SCS and CS to further our understanding of the evolution of this rare metabolic pathway and evaluate the pathways genesis. We include numerous archaeal and bacterial taxa, including some taxa not previously shown to have citrate-cleavage enzymes. Our results have several major implications for the evolution of autotrophy, but the abundance of horizontal gene transfer (HGTs) makes it uncertain whether the rTCA might have been active in LUCA.

## Methods

### Sequence retrieval

All amino acid sequences included in this study were obtained from NCBI GenBank database following BLAST searches of representative CCS, CCL, ACL, SCS and CS sequences in all major phyla of Bacteria and Archaea with a 30% identity cutoff. Enzyme identification of selected sequences was verified phylogenetically – that is, the largest monophyletic group that includes all examples of a respective enzyme (CCS, CCL, ACL, SCS or CS) previously characterized in the literature is labelled as these representative enzymes.

### Sequence analysis

Multiple amino acid sequence alignments were performed in NGPhylogeny software using MAFFT with default settings and BMGE curation. Tree inference was also conducted in NGPhylogeny with FastTree with LG and a gamma distribution for site-to-site rate heterogeneity, with 1000 bootstrap replicates and midpoint rooting (Lemoine et al., 2019). Phylogenies were visualized and edited in the Interactive Tree of Life (Letunic & Bork, 2021). We conducted an extensive review of properties of taxa included in this study using KEGG Pathway database (Kanehisa & Goto, 2000) and literature, cited in the Supplemental Table 1.

## Results and discussion

### Diversity of modern prokaryotes capable of citrate cleavage and rTCA

We conducted phylogenetic analysis with the citrate cleavage enzymes and their TCA homologs from a wide variety of bacterial and archaeal clades and examined the ecology of these clades. Our phylogenies support the homology between ACL and CCS/CCL complex first proposed by Aoshima, 2007 (Fig. 3, 4, 5). Additionally, recent advances in metagenomic studies and microbial systematics allowed us to identify and analyze significantly more potential homologs of these enzymes than done by Aoshima, 2007 and Beccerra et al., 2014. Particularly, we found for the first time, homologs of CCS in several groups of Acidobacteria, Deltaproteobacteria, Caldithricales and DPANN Archaea and homologs of ACL in Asgard Archaea, Bathyarchaeota and various bacterial clades (see Fig. 2 and Supplemental table 1). Some groups, such as Methanobacteria possess only a CCL enzyme, homologous to better characterized CCL of Aquificae and *Leptospirillum*.

**Fig. 2.**
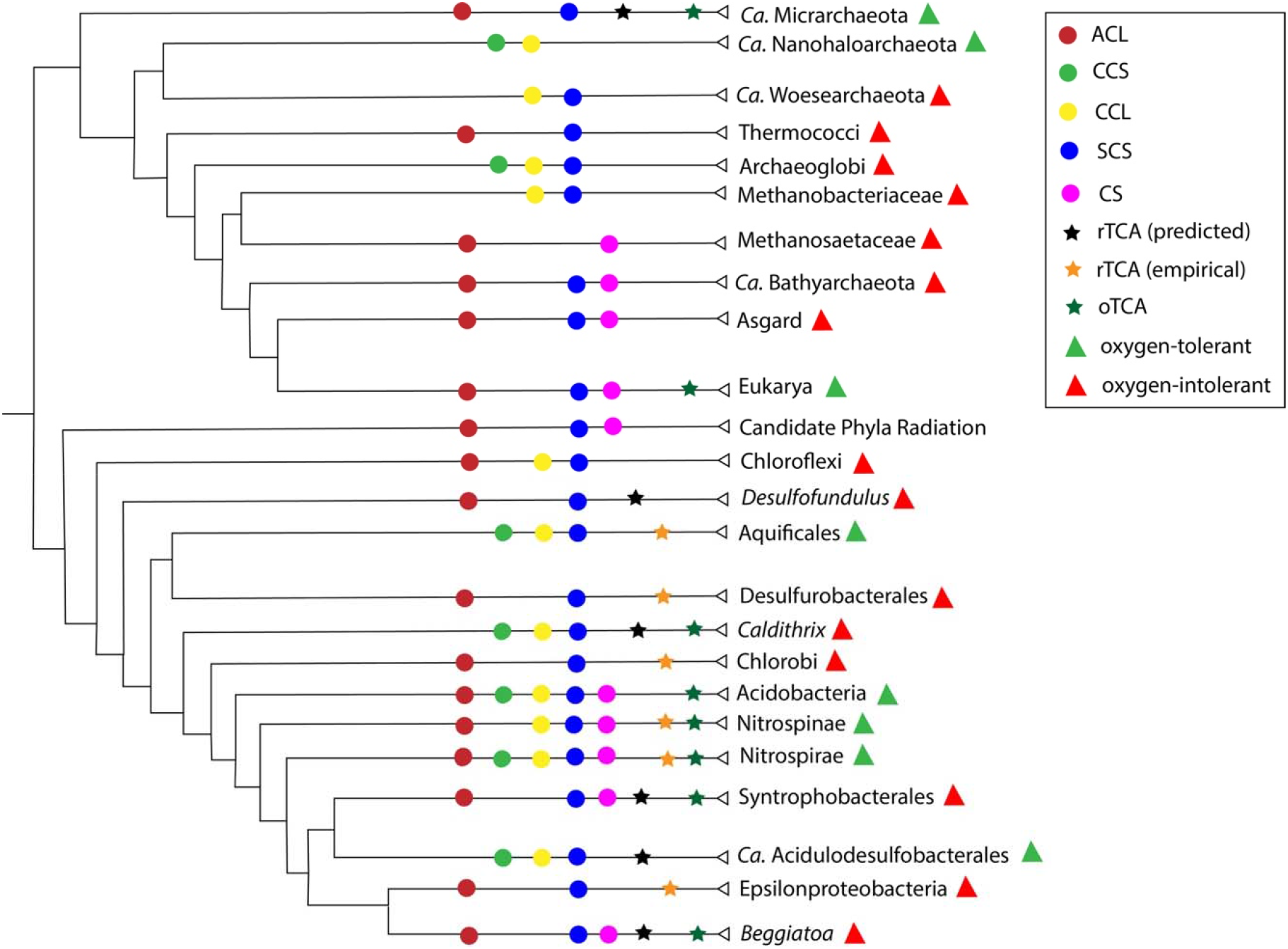
Phylogenetic distribution of citrate cleavage enzymes and associated traits in taxa included in this study. Tree topology is based on the tree of life phylogeny from Hug et al., 2016.

**Fig. 3.**
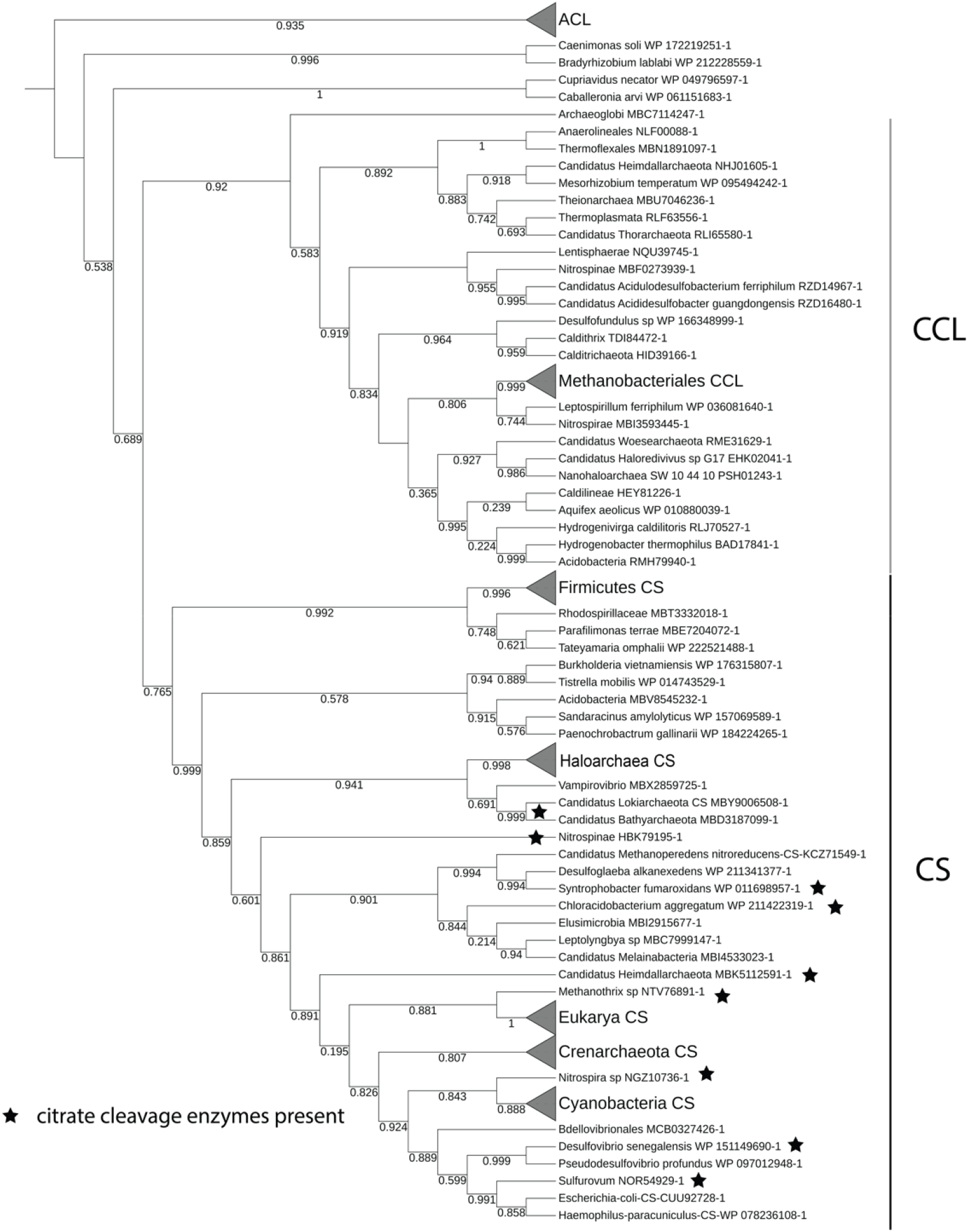
Phylogenetic tree of CS and CCL (maximum likelihood; 1000 bootstraps; bootstrap values labeled on the branches). Taxa that have representatives with both CS and citrate cleavage enzymes are marked with stars. Major monophyletic groups are collapsed for simplicity.

**Fig. 4.**
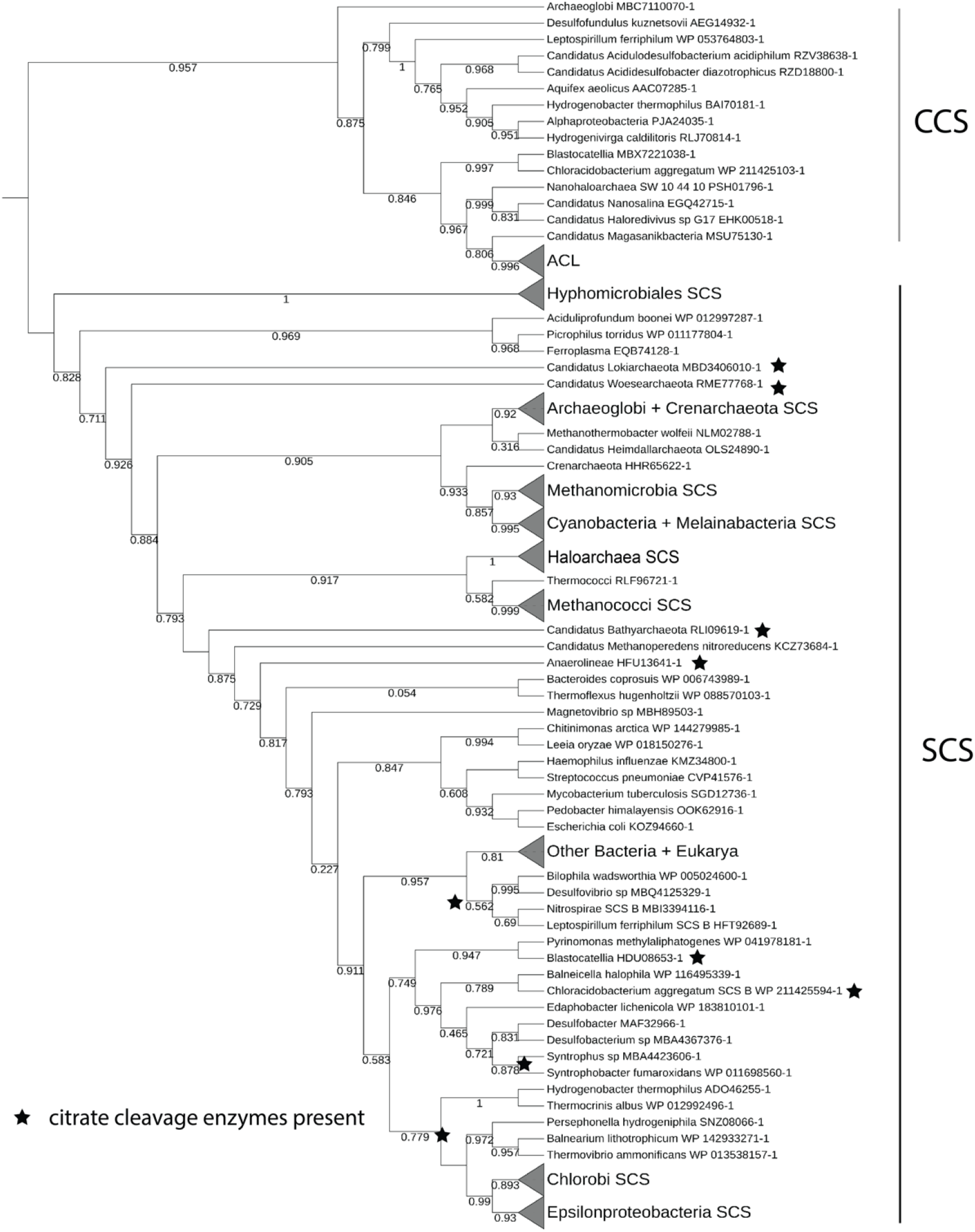
Phylogenetic tree of CCS and SCS β subunits (maximum likelihood; 1000 bootstraps; bootstrap values labeled on the branches). Taxa that have representatives with SCS and citrate cleavage enzymes are marked with stars. Major monophyletic groups are collapsed for simplicity.

**Fig. 5.**
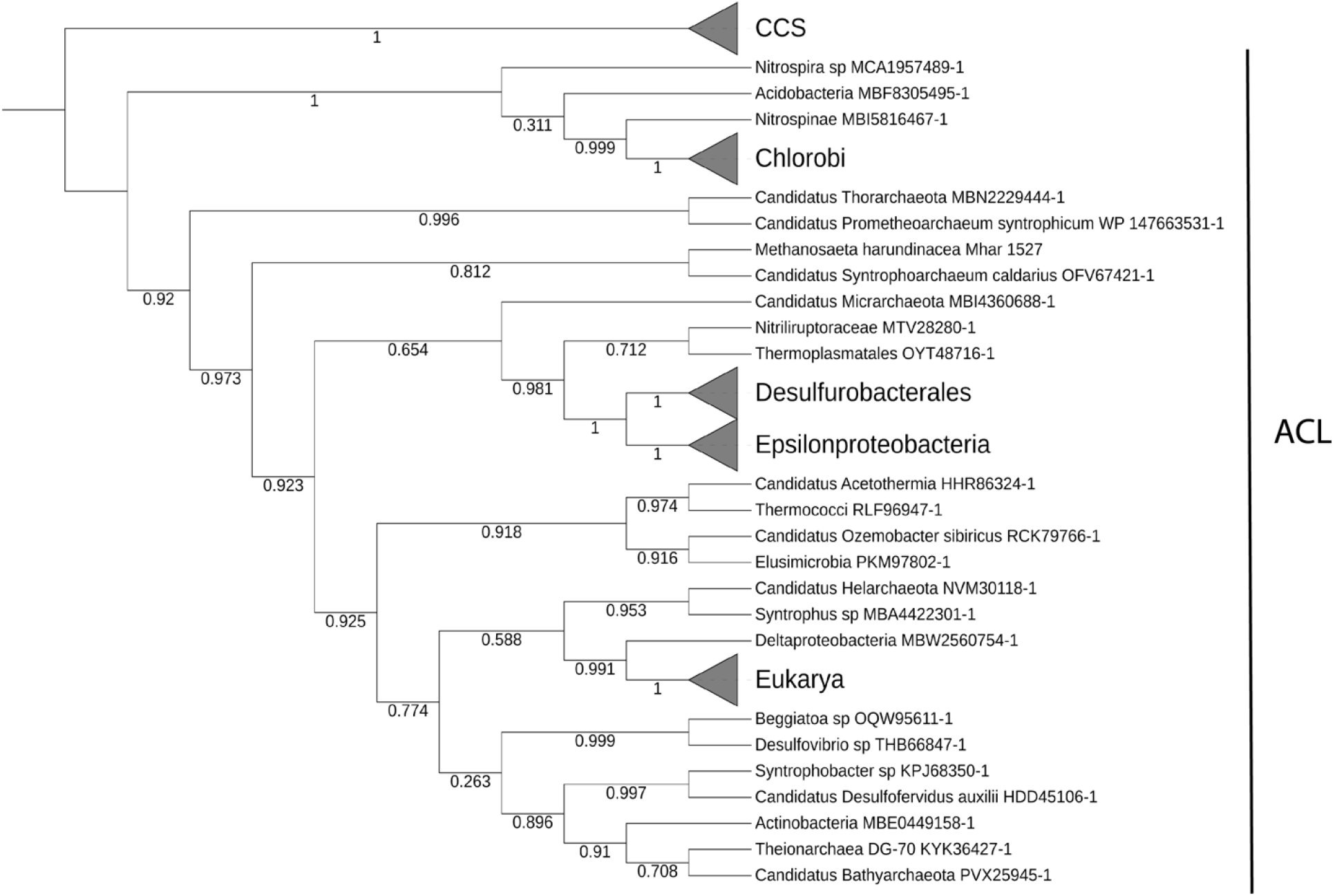
Phylogenetic tree of ACL β subunit (maximum likelihood; 1000 bootstraps; bootstrap values labeled on the branches). Major monophyletic groups are collapsed for simplicity.

Aoshima, 2007 proposed that the ACL enzyme resulted from a fusion between a CCS subunit and CCL, resulting in one-step catalysis of citrate cleavage into oxaloacetate and acetyl-CoA without releasing a citryl-CoA intermediate. Subsequently, two subunits of ACL – one derived from a fusion between CCS ß and CCL and the other from CCS α, fused again forming a one-subunit ACL enzyme (see Fig. 1, B). While the latter event likely only happened once, since a one-subunit ACL can only be found in eukaryotes (Chypre et al., 2012), the transition from CCS/CCL to ACL is less clear. These enzymes are generally poorly characterized, known mostly from environmental metagenomic studies and only a few type strains. CCS/CCL system is relatively well described only from Aquficales (Aoshima, 2004) and some members of Nitrospirae, such as *Leptospirillum* (Bar-Even et al., 2012), albeit our survey uncovered other homologies in DPANN Archaea (including Nanohaloarchaea), Methanobacteria, Theionarchaea, Deltaproteobacteria, *Caldithrix* and others, and multiple homologies for ACL enzymes. Few of these taxa contain all of the necessary TCA enzymes, however, we found several new taxa that could be theoretically capable of complete rTCA, at least based solely on bioinformatic evidence – specifically, Micrarchaeota, *Beggiatoa*, Ca. Acidulodesulfobacterales, *Syntrophobacter*, *Desulfofundulus* and Theionarchaea (see Supplemental Table 1 for an overview of enzyme presence in these taxa). Some of these enzymes are incorrectly annotated in GenBank as either SCS or CS despite being monophyletic with CCS or CCL respectively, as shown by our phylogenetic analyses. Specifically, these are sequences from Nanohaloarchaea and other DPANN Archaea and Candidate Phyla Radiation Bacteria (Fig. 3, 4). Our findings regarding distribution of taxa possessing citrate cleavage enzymes are summarized in Fig. 2.

Additionally, the studied citrate-cleaving taxa exhibit remarkable metabolic diversity (Fig. 2), contrary to what is usually suggested about modern rTCA distribution (Berg, 2011). Surprisingly, many CCS/CCL-possessing taxa were oxygen-tolerant (either aerobic or microaerophilic). A possible explanation for this could be due to easier regulation of oxidative and reductive TCA cycles when oxygen is present, since the CCS/CCL reaction releases citryl-CoA unlike the reverse CS reaction. This finding is inconsistent with the notion of rTCA representing an ancestral carbon fixation way, due to lack of oxygen on early Earth prior to the Great Oxygenation Event (Bar-Even et al., 2012). However, few of identified taxa in both CCS/CCL and ACL groups exhibit a complete rTCA cycle, hindering further examination of the connection between rTCA evolution and global oxygenation. An alternative plausible explanation for these findings is that during the CCS/CCL and ACL divergence, TCA was not cyclical in its modern form or that citrate cleavage reaction was involved in different ancient biochemical pathways. An additional complicating factor may have been the proposed horizontal transfer of aerobic respiration genes in some extant lineages (Kennedy et al., 2001; Ward et al., 2021).

### Patchy distribution of rTCA is due to HGTs, not multiple losses

To address the distribution of rTCA in detail, we examined phylogenetic distribution of SCS and CS – homologs of citrate-cleaving enzymes that also catalyze other TCA-related reactions. We did not find any meaningful patterns in the phylogeny of CS (Fig. 3), an enzyme that is generally unique to oTCA, and is also known to catalyze citrate cleavage with a ferredoxin cofactor in *Thermosulfidibacter takaii* (Nunoura et al., 2018) and *Hippea maritima* (Steffens et al., 2021). This is likely due to abundant HGTs in the evolutionary history of this enzyme. On the other hand, homologs SCS and CCS catalyze relatively simple thermodynamically favorable reactions (for succinyl-CoA synthesis ΔG°= −1.2 kJ/mol and for citryl-CoA synthesis ΔG°= −5.9 kJ/mol according to eQuilibrator web server, Flamholz et al., 2012) and both reactions are necessary for a complete rTCA cycle. Interestingly, in our phylogeny, SCS from most otherwise unrelated Bacteria capable of citrate cleavage with either CCS/CCL or ACL form a monophyletic group (Fig. 4). These groups include Aquificae, Chlorobi, Epsilonproteobacteria and various Deltaproteobacteria and Acidobacteria in which we found homologs of citrate-cleavage enzymes. This finding can be interpreted by Bacteria obtaining their SCS, CCS and ACL from a single linked HGT event since genes for these and other TCA-related enzymes are often located in close proximity to each other in the genomes (according to our searches of these enzymes in the SynTax database, Oberto, 2013). These results are congruent with the explanation of patchy distribution of rTCA through abundant horizontal gene transfers rather than multiple gene loss events, as was previously suggested (Aoshima, 2004).

The deepest evolutionary relationships of these enzymes are difficult to elucidate with bioinformatical tools alone. However, our ecological and phylogenetic comparisons described above, seem to broadly indicate that a complete rTCA cycle in modern taxa might be a relatively recent evolutionary innovation due to abundant HGTs. We will also discuss a possible alternative later in this section. Since SCS is also involved in oTCA, the possibility of the complete TCA cycle being a late evolutionary innovation in either direction is certainly worthy of further investigation. In fact, a connection between evolution of TCA and relatively late events such as eukaryogenesis or Great Oxygenation Event was previously raised by Ryan et al., 2021. Since we found ACL in multiple clades of Asgard Archaea, it is logical to assume that eukaryotic ACL is derived from their Asgard ancestors rather than an alphaproteobacterial mitochondrial endosymbiont. Significantly, however, amino acid sequences from eukaryotic ACL proteins do not form a monophyletic group with their Asgard homologs (Fig. 5), potentially implicating further unidentified HGT events either in Eukarya or stem Asgard Archaea.

### Cyanobacterial SCS originated from an archaeal HGT

A major implication of our findings is that the evolution of enzymes catalyzing different steps of the TCA cycle – a ubiquitous metabolic pathway in modern life, was riddled with horizontal gene transfers. Particularly with SCS, the phylogenetic analysis suggests that Cyanobacteria/Melainabacteria clade obtained their SCS from a group of thermophilic Archaea, possibly Methanomicrobia, through HGT (Fig. 4). Cyanobacteria belonging to various orders and Melainabacteria make a monophyletic group with two euryarchaeote groups - Methanomicrobia and Archaeoglobi and Crenarchaeota. These three major archaeal clades occupy distant positions in most phylogenetic analyses of Archaea (Matte-Tailliez et al., 2002). However, they share similar ecologies, largely being thermophilic. A notable exception is Methanomicrobia – some representatives are only moderately thermophilic (Battumur et al., 2019), while some genera, such as *Methanosaeta* and *Methanothrix* lack the SCS enzyme. Interestingly, this is not the only example of an HGT event from Archaea to Cyanobacteria. For instance, structural maintenance of chromosomes (SMC) genes were shown to be transferred from methanogens (Soppa, 2001). SMC transfer event was used to constrain the evolutionary timeline for Cyanobacteria and methanogenic Archaea (Wolfe & Fourier, 2018), while the HGT of SCS enzymes could be potentially used to resolve cyanobacterial evolution even further since it occurred prior to Cyanobacteria/Melainabacteria divergence. More studies on TCA-related genes are necessary to identify a specific donor group for the SCS gene among methanogenic Archaea.

The divergence between Cyanobacteria and their non-photosynthetic relatives Melainabacteria was found to predate the Great Oxygenation Event, happening approximately 2.5 billion years ago (Shih et al., 2017). In addition to being anaerobic, Melainabacteria are distinct in multiple ways from Cyanobacteria, sometimes included in the class Oxyphotobacteria, due to their capacity for oxygenic photosynthesis (Di Rienzi et al., 2013). Most Melainabacteria possess only complex I of the electron transport chain (ETC) that can function in anaerobic environments, implying that aerobic ETC post-dated the great oxygenation (Grettenberger et al., 2021). However, some groups of Melainabacteria independently acquired aerobic respiratory complexes, such as Vampirovibrionales (Soo et al., 2017).

An interesting common feature of all of these clades – thermophilic Archaea, Cyanobacteria and Melainabacteria is the incompleteness of their TCA cycles (Slobodkina et al., 2021). Modern Cyanobacteria lack oxoglutarate dehydrogenase (OGDH), a key TCA enzyme that converts 2-oxoglutarate into succinyl-CoA, and instead utilize a succinic semialdehyde shunt (Steinhauser et al., 2012). A possible explanation for this unique version of the TCA cycle is its connection to nitrogen metabolism through succinic semialdehyde amination to produce gamma aminobutyric acid (Zhang et al., 2016). Additionally, replacing succinyl-CoA with succinic semialdehyde as a TCA cycle intermediate decreases cumulative free energy and therefore increases forward flux through the cycle (Thomas et al., 2014). While the succinic semialdehyde shunt bypasses succinyl-CoA as an intermediate in the TCA cycle, SCS is still retained in Cyanobacteria likely due to succinyl-CoA being one of the key biochemical precursors, being necessary for synthesis of several amino acids. Melainabacteria and Archaeoglobi lack most of the TCA enzymes, including OGDH and there is no evidence of succinic semialdehyde being produced instead (Di Rienzi et al., 2013). Furthermore, Crenarchaeota lack OGDH but instead use a ferredoxin-dependent enzyme to catalyze succinyl-CoA synthesis (Hallam et al., 2006).

In addition to phylogeny, we also compared the citrate-binding region of CCS B subunit, obtained from the X-ray diffraction structure of *Hydrogenobacter thermophilus* CCS (PDB ID: 6HXQ; Verschueren et al., 2019), to various ACL and SCS proteins. This region consists of eight amino acids, all in the beta subunit: F270, G296, G297, A327, N328, N329, T330, R363. This region exhibits 50% similarity to the respective aligned sequence of *Chlorobium* and *Methanosaeta* ACL enzymes and only 25% to SCS proteins from all of the studied taxa, with only two consecutive glycines remaining conserved. Although, among bacterial, archaeal and eukaryotic SCS proteins this region is highly conserved as well with variation only in two positions. The only exceptions are cyanobacterial SCS sequences, which are different from both CCS and other SCS proteins in almost every position, suggesting either erroneous homology or significant functional divergence. For example, SCS in Cyanobacteria might be conserved not for the TCA but due to the importance of succinyl-CoA in amino acid biosynthesis.

### Possibility of rTCA operation within syntrophic consortia

Out of the taxa included in this study, Haloarchaea and their symbiotic counterparts, Nanohaloarchaea, are some of the most interesting. Both groups are adapted to highly saline environments, however Nanohaloarchaea exhibit significant genomic and metabolic reduction. Phylogenetic relationships of Nanohaloarchaea are also unclear, with their placement in DPANN super-phylum being challenged by Aouad et al., 2018. On the other hand, Haloarchaea are frequently suggested to have evolved from a lineage of methanogenic Archaea (Nelson-Sathi et al., 2015). Modern methanogens often couple methanogenesis to carbon fixation pathways, specifically Wood- Ljundahl pathway (Borrell et al., 2016), however Haloarchaea are exclusively heterotrophic and have a phototrophic capability. In addition, they possess bacteriorhodopsins – light-harvesting proton pumps, that unlike photosystems in Cyanobacteria and eukaryotes are not implicated in any redox processes. In fact, it was suggested by DasSarma & Schwieterman, 2021 that chlorophyllic photosynthesis originated due to competition for wavelength with retinal-based photosystems – a Purple Earth hypothesis.

A major issue with this hypothesis is that Haloarchaea, one of the most prominent modern groups that use retinal-based proton pumps, are not known to fix carbon and couple the resulting proton gradients to reduction of electron carriers, unlike, for example, Cyanobacteria with Calvin-Benson cycle or Chlorobi with rTCA. While there is no evidence of bacteriorhodopsin involvement in redox processes, one carbon fixation pathway in Haloarchaea is almost complete – the rTCA. Haloarchaea possess all necessary enzymes for rTCA with the exception of an enzyme for citrate cleavage. Haloarchaea are also known to engage in intimate symbioses with Nanohaloarchaea, as was recently shown for the relationship between *Candidatus Nanohalobium* and *Halomicrobium* (La Cono et al., 2020) and *Candidatus Nanohaloarchaeum* and *Halorubrum* (Hamm et al., 2019). Out of all rTCA enzymes we found Nanohaloarchaea, including *Candidatus Nanohalobium* and *Candidatus Nanohaloarchaeum*, to possess only two: both CCS and CCL, indicating these organisms are capable of citrate cleavage. This raises the tantalizing possibility that Haloarchaea and Nanohaloarchaea may, together, be able to couple retinal-based proton pumping to carbon fixation via the rTCA (Fig. 6). Along similar lines, Methanomicrobia which often lack ACL, are known to engage in syntrophic relationships with at least one Asgard Archaeon, *Prometheoarchaeum syntrophicum* (Imachi et al., 2020). Since we found that most Asgard phyla possess ACL, the syntrophic consortium might be able to perform the complete rTCA. Likewise, Deltaproteobacteria from Syntrophobacterales possess ACL and are involved in hydrogen syntrophy with archaeal methanogens (Harmsen et al., 1998). Albeit, these findings require further *in vivo* investigations.

**Fig. 6.**
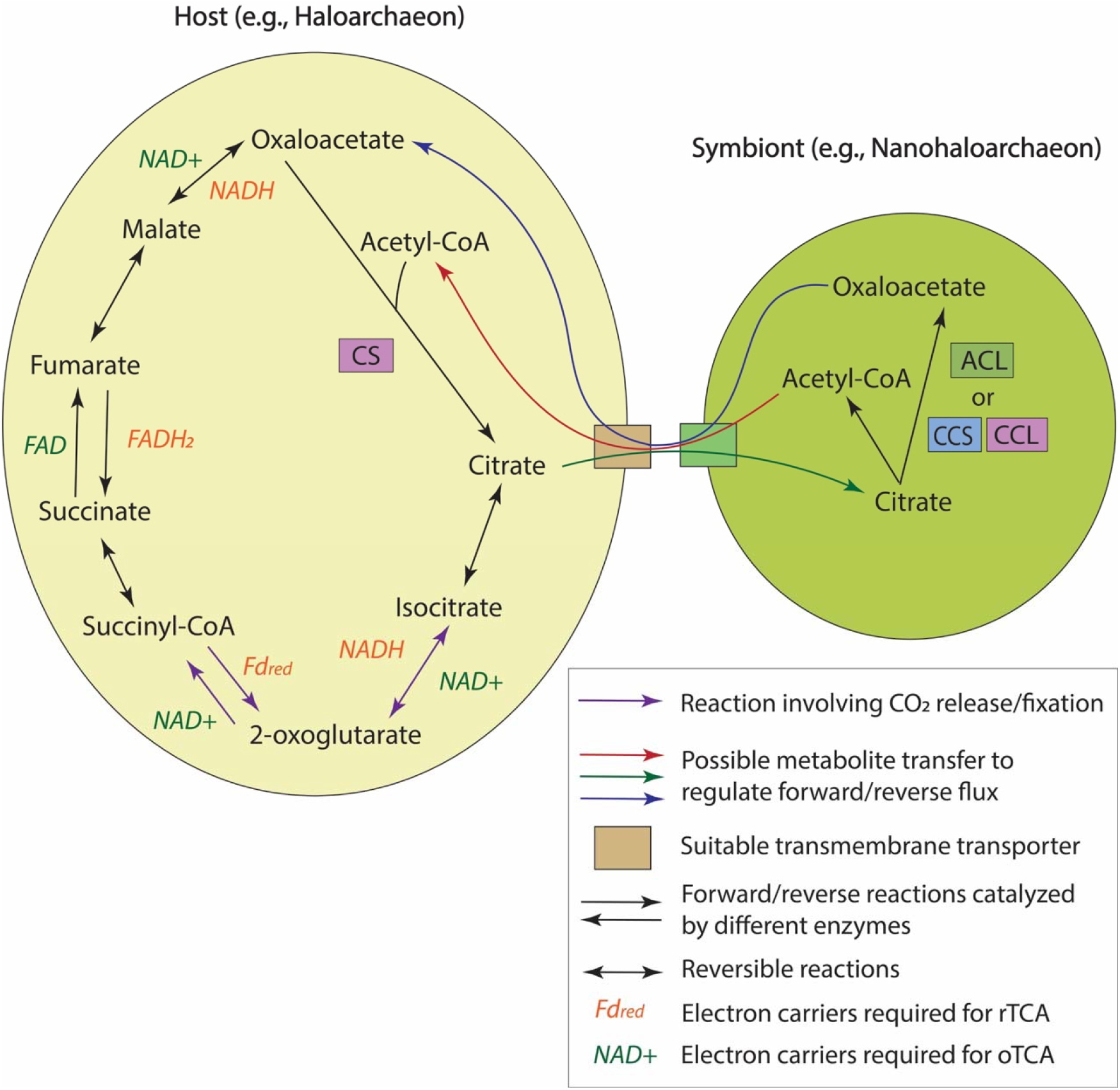
Schematic depiction of possible syntrophic operation of the TCA cycles between a host and a symbiont, such as a Haloarchaeon and Nanohaloarchaeon (Hamm et al., 2019; La Cono et al., 2020). Electron carriers required in either oxidized or reduced form for either oTCA or rTCA reactions are noted (Fd_red_ – reduced form of ferredoxin).

Inter-symbiont gene transfer is known for another archaeal association – a Crenarchaeote *Ignicoccus hospitalis* and a member of the DPANN superphylum *Nanoarchaeum equitans* (Podar et al., 2008). Multiple studies of this association have shown consistent protein and metabolite transfer between the two partners to compensate for shortcomings of their individual metabolomes (Giannone et al., 2011), showing that a similar system is theoretically possible for a Nanohaloarchaeon and a Haloarchaeon, or a member of Methanomicrobia with its bacterial or archaeal symbiotic partner. Both Nanohaloarchaea and Haloarchaea possess copies of genes of various transporters capable of moving small metabolites across the membrane, such as the major facilitator superfamily transporters (Quistgaard et al., 2016).

### Syntrophic separation of oxidative and reductive fluxes through the TCA cycle

Inter-symbiont transfers of enzymes or metabolites can form a basis for many syntrophic associations, however, CO_2_-fixing reactions in particular are very thermodynamically challenging, and may require over-reduction of electron carriers or switching to lower reduction carriers, such as ferredoxins, implicated in rTCA (Bar-Even et al., 2012). An excess of electrons can be hard to come by without the presence of multiple specialized oxidative pathways. Plants and other organisms using Calvin-Benson cycle possess light-activated photosystems capable of reducing NADP, while most Archaea lack systems of comparable efficiency. Hence, different groups of organisms use different strategies to regulate fluxes through TCA reactions, for example, with citrate synthase capable of catalyzing citrate cleavage dependent on CO_2_ availability in *Hippea maritima* (Steffens et al., 2021) and *Thermosulfidibacter takaii* (Nunoura et al., 2018). However, splitting the operation of rTCA cycle into two syntrophic cells might be beneficial to allow for regulation of both forward and reverse fluxes through the cycle with lesser dependence on environmental conditions or availability of reduced electron carriers, which can be either produced in oxidative reactions or consumed in the reductive ones (Fig. 6). Citrate cleavage reaction is a particularly good candidate for such syntrophic regulation since it is far less energetically expensive than direct CO_2_ fixation reactions.

This brings us back to Haloarchaea. Despite not providing direct electron sources, retinal-based photosystems do effectively supply other sympatric microorganisms with proton gradients and could be viewed as an efficient tool for a mutualist. Moreover, regarding their ancient origins and possibly wider distribution (DasSarma & Schwieterman, 2021), it is plausible that ancestral retinal-phototrophic Archaea could have obtained necessary electrons through syntrophy with microorganisms possessing more diverse oxidative pathways, such as sulfide or nitrite oxidation, metal oxidation or even water splitting. The syntrophic electron transfer process itself is a relatively conserved process, that could occur in several ways – either through multi-heme cytochrome c networks, reduction-oxidation of extracellular Fe/Mn (Shi et al., 2016) or hydrogen/formate generation (Nobu et al., 2014). Multi-heme cytochromes c are absent in Haloarchaea, but present in related methanogens (Kletzin et al., 2015). Additionally, it seems that Haloarchaea lack any efficient methods of metal oxidation. However, they possess a formate dehydrogenase and interspecies formate transfer is a common feature of syntrophic relationships of methanogens (Hattori et al., 2001). For example, this is a proposed method of syntrophy for methanogens aggregating with a recently cultured *Prometheoarchaeum syntrophicum* (Imachi et al., 2020).

Role of syntrophy and symbioses in the early evolution of life is frequently overlooked. In fact, such ecosystemic approach can be extended to the origins of life as a whole, suggesting a heterogeneous population of proto-organisms prior to LUCA with high levels of metabolic specialization and reliance on obligate symbioses. This is essentially a continuation of the theory of autonomous functional systems suggesting heterogenous molecular populations in the early prebiotic chemistry (Ruiz-Mirazo et al., 2017) and synergistic selection theory implying the importance of synergy in the early evolution of life (Corning & Szathmary, 2015). It was also demonstrated that syntrophy emerges spontaneously within complex metabolic networks (Libby et al., 2019) and that syntrophy in modern microbes is significantly more common than anticipated (Morris et al., 2013). This idea would be interesting to explore further, because it is supported by several features of life on Earth, such as abundance of nano-prokaryotes (DPANN and Candidate Phyla Radiation clades), late origins of intracellular membranes, or abundance of Fe-S proteins and Ni-Fe hydrogenases in LUCA (Sutherland, 2017).

### Conclusions

In this study we aimed to understand the evolution of reductive TCA cycle, a carbon fixation pathway of major significance to the field of origins of life studies. We show that the strangely sporadic distribution of this pathway in modern microorganisms can be largely explained by abundant horizontal gene transfers. Additionally, we suggest that evolution of rTCA could have been determined by interactions within syntrophic consortia to increase metabolic flexibility and ease switching between oxidative and reductive fluxes. We also highlight the importance of an ecological perspective to the question of the origins of life as a whole.

## Abbreviations

rTCA: reductive Tricarboxylic Acid cycle
oTCA: oxidative Tricarboxylic Acid cycle
LUCA: Last Universal Common Ancestor
ACL: ATP-citrate lyase
CCS: citryl-CoA synthase
CCL: citryl-CoA lyase
SCS: succinyl-CoA synthase
CS: citrate synthase
HGT: horizontal gene transfer

## Acknowledgements

This research was conducted during Blue Marble Space Institute of Science Young Scientist Program (YSP) in 2021. The authors would like to thank Priya DasSarma for assistance with phylogenetic analysis and members of the DasSarma YSP 2021 group, David Baum and Praful Gagrani for providing helpful feedback on this study and the manuscript. The DasSarma laboratory is supported by NASA Exobiology grant 80NSSC17K0263.

