## Supplemental Table 1 for "Role of horizontal gene transfers and microbial ecology in the evolution of fluxes through the tricarboxylic acid cycle"

**Supplementary Table 1.** Summary of taxa found to possess citrate cleavage enzymes and their metabolic properties. This data is based on available genomic databases, in which case a unique identifier is provided in the table, or on the literature that is cited in the table and referenced below.
